# Reverse transcription quantitative PCR to detect low density malaria infections

**DOI:** 10.1101/2020.07.01.183491

**Authors:** Peter Christensen, Zbynek Bozdech, Wanitda Watthanaworawit, Laurent Renia, Benoit Malleret, Clare Ling, Francois Nosten

**Affiliations:** Shoklo Malaria Research Unit, Mahidol-Oxford Tropical Medicine Research Unit, Faculty of Tropical Medicine, Mahidol University, Maesot, Thailand; Department of Microbiology and Immunology, University of Otago, Dunedin, New Zealand; School of Biological Sciences, Nanyang Technological University, Singapore, 637551; Singapore Immunology Network, A*STAR, Singapore, 138648; Department of Microbiology and Immunology, Yong Loo Lin School of Medicine, National University of Singapore, Singapore, Republic of Singapore; Centre for Tropical Medicine and Global Health, Nuffield Department of Medicine, University of Oxford, Oxford, United Kingdom

## Abstract

Targeted malaria elimination strategies require highly sensitive tests to detect low density malaria infections (LDMI). Commonly used methods for malaria diagnosis such as light microscopy and antigen-based rapid diagnostic tests (RDTs) are not sensitive enough for reliable identification of infections with parasitaemia below 200 parasites per milliliter of blood. While targeted malaria elimination efforts on the Thailand-Myanmar border have successfully used high sample volume ultrasensitive quantitative PCR (uPCR) to determine malaria prevalence, the necessity for venous collection and processing of large quantities of patient blood limits the widespread tractability of this method. Here we evaluated a real-time quantitative reverse transcription PCR (qRT-PCR) method that significantly reduces the required sample volume compared to uPCR. To do this, 304 samples collected from an active case detection program in Kayin state, Myanmar were compared using uPCR and qRT-PCR. *Plasmodium* spp. qRT-PCR confirmed 18 of 21 uPCR *Plasmodium falciparum* positives, while *P. falciparum* specific qRT-PCR confirmed 17 of the 21 uPCR *P. falciparum* positives. Combining both qRT-PCR results increased the sensitivity to 100% and specificity was 95.1%. These results show that malaria detection in areas of low transmission and LDMI, can benefit from the increased sensitivity of qRT-PCR especially where sample volume is limited.

## Background

As malaria burden reduces globally, the international community is working toward its elimination. Successful targeted malaria elimination strategies will require increased surveillance and highly sensitive tests capable of detecting asymptomatic and low density malaria infection (LDMI). These infections are often well below 200 parasites per milliliter and are an important disease reservoir capable of transmitting malaria that must be detected and eliminated (1, 2). Light microscopy and antigen based rapid diagnostic tests (RDTs) are the most common tests used in malaria prevalence surveys, and usually assess 5 μL of whole blood per test, a volume which precludes reliable detection of LDMI. Ultrasensitive RDTs improve detection sensitivity of patients with a parasitaemia between 200 parasites/mL and 10,000 parasites/mL (3), but are still limited by their low input volume. Molecular methods using the polymerase chain reaction (PCR) remain the only common and reliable method to detect LDMI. This technique’s sensitivity is due to its ability to detect single, specific DNA molecules and use concentrated DNA from a large sample. Widespread use of PCR based assays, namely real-time quantitative PCR (qPCR) and reverse-transcription qPCR (qRT-PCR), have revealed a new landscape of malaria prevalence particularly in low transmission areas (4, 5).

The targeted malaria elimination project (TME) on the Eastern Myanmar border used a high blood volume ultrasensitive qPCR (uPCR) to consistently detect parasitaemia down to 22 parasites per mL (6), and revealed a high proportion of LDMI (7). Major features of uPCR are its 7 copies of gene target, the high volume of blood tested and the ability to accurately quantify low density parasitaemia. Although increasing the blood volume of a PCR leads to higher sensitivity (6), the collection of large numbers of high volume samples have their own specific limitations. These can include, the ethics approval required for venous blood draw, sample logistics, increased nucleic acid extraction cost and increased sample processing time. Another way to increase the sensitivity of PCR is to increase the number of specific nucleic acid targets per parasite by targeting specific RNA and DNA using qRT-PCR. As previously reported by Kamau et al. (8), the primer set used in uPCR can be made more sensitive by incorporating reverse transcription prior to qPCR, enabling detection of the 7 genes encoding Plasmodium 18S ribosomal nucleic acid (rRNA) and its rRNA transcripts.

In this study, we compare the sensitivity and specificity of high sample volume, low target copy number ultrasensitive qPCRs, with reduced sample volume, high target copy number qRT-PCR for the detection and quantification of *Plasmodium* spp. and *P. falciparum.*

## Methods

We selected 304 samples with previously reported uPCR results: 21 *P*. *falciparum* positive and 283 *Plasmodium* spp. negative for comparison to qRT-PCR with increased target numbers per parasite but 30% of the sample volume.

### Study area and sample collection

Active case detection samples were collected from rural Eastern Kayin (Karen) state of Myanmar between 2014 and 2015 as part of an international malaria elimination project. Full methods have been published (9), briefly, 3 ml of blood was drawn into an EDTA container from each adult, and transported on ice to the Shoklo Malaria Research Unit in Mae Sot, Tak, Thailand. Within 48 hours the samples were processed, and two aliquots of up to 500 μL of packed red blood cells (PRBC) were stored at −80°C. Only 500 μL samples were accepted for METF qPCR while qRT-PCR included samples with at least 150 μL.

### Ethics Statement

First, community engagement teams sought community approval ahead of the survey date. Then, survey participants received individual information in their language, and informed consent was obtained from each individual before they provided a venous blood sample. Appropriate treatment for *Plasmodium falciparum* or *Plasmodium vivax* was available for all RDT-positive individuals.

The METF project has ethical approval from the Lower Myanmar Department of Medical Research Ethics’ committee (reference 73/ETHICS2014, dated 25 November 2014, and renewed in November 2015 and 2016 under the same reference).

### Ultrasensitive qPCR (uPCR)

DNA was extracted from 500 μl of cryopreserved PRBC using QIAamp DNA Blood Midi Kit according to manufacturer’s instructions. The DNA template was then dried in a vacuum concentrator, re-suspended in 10 μL of PCR grade water and stored at −20°C prior to qPCR. Separate uPCRs specific for *Plasmodium* spp., *P. falciparum* and *P. vivax* were performed over 3 years from 2013 as previously described (6). Each 10 μL uPCR reaction contained 2 μL of DNA template with 1x QuantiTect Multiplex PCR No ROX mastermix (Qiagen^™^), 0.4 μM each primer, and 0.2 μM Taqman probe. Thermal cycling and signal acquisition was done on an ABI 7500 Fast real-time PCR machine with initial denaturation and enzyme activation at 95°C for 15 min, then 50 cycles of denaturation at 94°C for 15 sec followed by annealing and extension at 60°C for 60 sec. A reaction with exponential signal increase before cycle 40 was considered positive.

### Sample selection, Nucleic acid extraction and qRT-PCR

Within 3 years of sample storage at −80°C, nucleic acid extraction and qRT-PCR assays were done on the second aliquot of PRBC for selected samples: 21 uPCR *P. falciparum* positives and 283 uPCR negatives.

Nucleic acid was extracted using Quick-RNA Miniprep (Plus) kits from Zymo Research. Manufacturer’s instructions for whole blood were followed with minor changes. These include, extraction from 150 μL of white blood cell depleted PRBC in phosphate buffered saline (PBS) up to 200 μL instead of 200 μl of whole blood, DNase enzyme wasn’t used, and RNA was eluted in 20 μL of molecular grade water. Two qRT-PCRs were done on each extract in duplicate: the *Plasmodium* spp. specific assay using the same primer and probe set as uPCR, and a *P. falciparum* specific set targeting the DNA of the A-18S rRNA genes and its rRNA transcripts (5). Both reactions had a final volume of 15 μL and contained 4 μL of RNA template, 1x Superscript III One-Step RT-PCR System master mix (ThermoFisher Scientific^™^), 0.4 μM forward and reverse oligo primer and 0.2 μM of MGB Taqman probe. Amplification and signal acquisition were done on an ABI 7500 Fast real-time PCR machine with cycling conditions as follows: reverse transcription at 45°C for 30 min, enzyme activation at 95°C for 2 min, followed by 50 cycles of denaturation at 95°C for 15 sec and combined annealing and extension steps at 60°C for 60 sec. A reaction with exponential signal increase before cycle 40 was considered positive.

### Standard Curve

Standard reference curves for the qRT-PCR and uPCR were made using aliquots of 10,000 flow cytometry sorted small ring stage *P. falciparum* (3D7) parasites (10). The method used to extract the nucleic acids from these parasites depended on the assay used (qRT-PCR or uPCR). For the uPCR standard curve, Qiagen DNA Blood Mini Kit was used to extract DNA, this was eluted in 200 μL of sterile water, dried in a partial vacuum at 30°C for consistency with sample extraction, and re-suspended in 200 μL of Qiagen AE buffer (6). Nucleic acid for the qRT-PCR standard curve was extracted using the Zymo Quick-RNA Miniprep (Plus) kit as above but eluted in 40 μL of water. Serial one in five dilutions were made with these extracts to make 7 standards. The uPCR standard curve ranged from 100 to 0.032 parasites per qPCR reaction and the qRT-PCR standard curve ranged from 1000 parasites to 0.064 parasites per reaction.

### Analysis

*Plasmodium* spp and *P. falciparum* qRT-PCR results were analysed using 2 x 2 contingency tables with uPCR as gold standard. Results of both qRT-PCRs were also combined and compared to uPCR, where a positive test by at least one qRT-PCR was considered positive.

## Results

### *Plasmodium* spp. qRT-PCR

*Plasmodium* spp. qRT-PCR confirmed 18 of 21 *Plasmodium* spp. uPCR positives and an additional 9 positive reactions from the 283 uPCR negatives. The sensitivity and specificity of *Plasmodium* spp. qRT-PCR was 85.7% and 96.8% respectively when compared to uPCR (Table 1). The positive predictive value (PPV) and negative predictive value (NPV) of this test was 66.7% and 98.9% respectively.

**Table 1.** Results of three PCRs on 304 malaria survey samples: tabulated results from *Plasmodium spp.* uPCR, *Plasmodium spp.* RT-qPCR and *P. falciparum* RT-qPCR including total positives, total negatives, false positives, false negatives, sensitivity and specificity using *Plasmodium spp.* uPCR results as gold standard.

|  | Positive | Negative | False Positive | False Negative | Sensitivity | Specificity |
| --- | --- | --- | --- | --- | --- | --- |
| *Plasmodium* spp. uPCR | 21 | 283 | N/A | N/A | N/A | N/A |
| *Plasmodium* spp. qRT-PCR | 27 | 277 | 9 | 3 | 85.7% | 96.8% |
| *P. falciparum* qRT-PCR | 21 | 283 | 5 | 5 | 76.2% | 98.2% |
| Combined qRT-PCR | 35 | 269 | 14 | 0 | 100% | 95.1% |

### *P. falciparum* qRT-PCR

Using the *P. falciparum* specific qRT-PCR, 16 of the 21 uPCR *P. falciparum* positives were confirmed along with 5 extra positives from the 283 negatives. Sensitivity and specificity of this test was 76.2% and 98.2% respectively when compared to uPCR (Table 1). PPV for this PCR was 76.2% and NPV was 98.2%.

### Combined *Plasmodium* spp. and *P. falciparum* qRT-PCR

Combining the results of both *Plasmodium* spp. and *P. falciparum* qRT-PCRs, all 21 uPCR positives are confirmed positive (100% Sensitivity) with 14 false positives (specificity 95.1%) (Table 1). These combined results gave a PPV of 60% and a NPV of 100%.

### Quantification

The quantification by each method gave highly varied results. Repeated measures one way analysis of variance with post hoc analysis using Tukey’s multiple comparison test revealed no significance between parasitaemia counts of each PCR technique (Figure 1).

**Figure 1.**
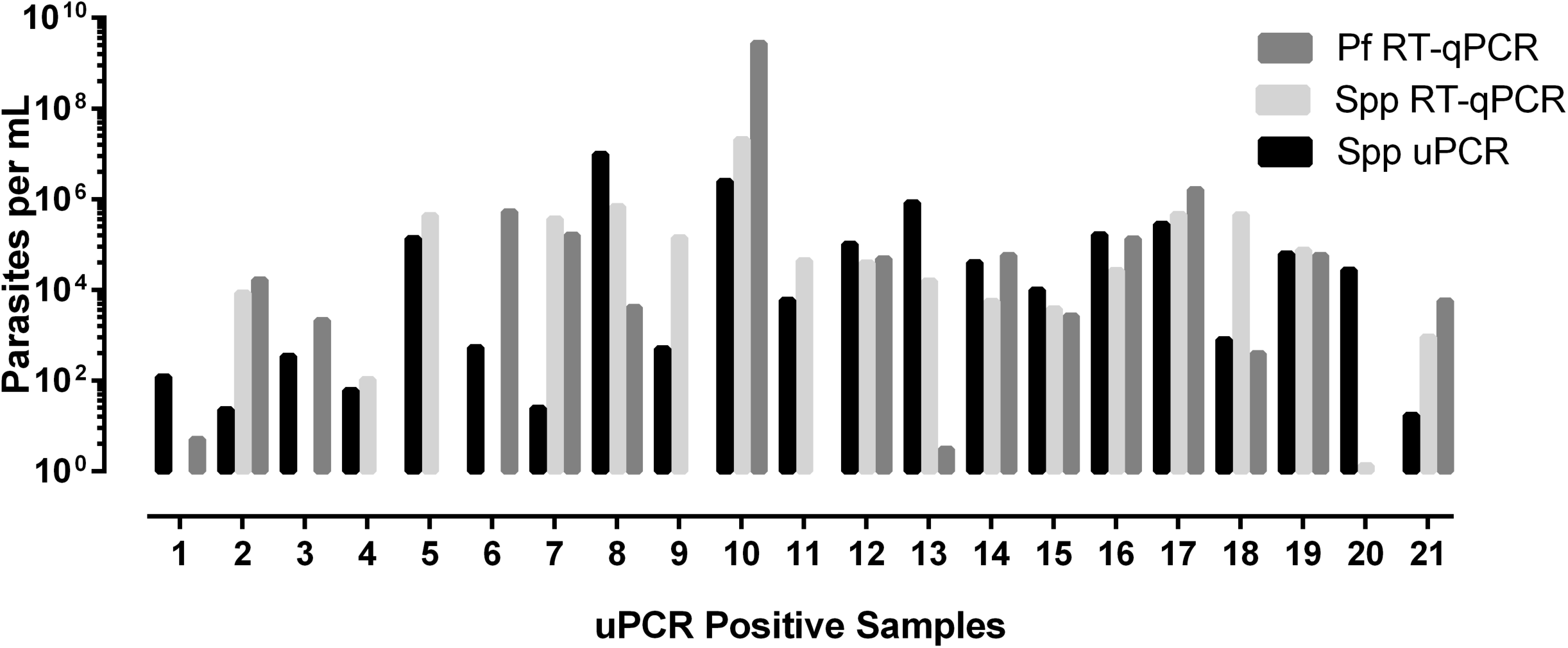
Parasite quantification by spp. uPCR, spp. RT-qPCR and Pf RT-qPCR. Quantification of parasitaemia from three PCRs on 21 uPCR positives from the TME malaria survey of Kayin state, Myanmar. Parasitaemia by uPCR ranged from 17 to 9.91×10^6^ parasites per mL whole blood. One-way ANOVA with multiple comparisons revealed no significance between quantification results.

## Discussion

LDMI detection is essential for effective targeted elimination programs, necessitating the careful selection of a detection assay that is appropriate to the setting and study requirements. Sample volume, storage conditions and transit time are important factors, as well as the required assay sensitivity, specificity and its cost. RNA is generally less stable than DNA, and accurate RNA quantification often requires normalization due to variable transcription rates. It is for these reasons that DNA based qPCR was chosen for the original study as this approach enabled accurate quantification of LDMI in a setting where samples from remote locations would likely experience delays.

The targeted malaria elimination (TME) project on the Eastern Myanmar border used conserved regions of the 18S rRNA genes as the target for qPCR. This high sample volume uPCR assay was modified from the qRT-PCR published by Kamau et al. 2011 for their detection of low density malaria infections (LDMI). After uPCR detected a high prevalence of LDMI in the region (9), and with continued surveillance in mind, we wanted a lower sample volume assay with similar LDMI detection sensitivity. As Kamau et al. has shown, using this primer set as a qRT-PCR significantly increases the sensitivity of the assay. The increased sensitivity of qRT-PCR is due to the increased number of targets per parasite (compared to uPCR). These amplification targets include the 18S rRNA genes, and the structural RNA of each ribosome. *Plasmodium* genus specific uPCR amplifies type A and S 18S ribosomal RNA genes distributed on chromosomes 1, 5, 7, 11 and 13. A positive qPCR reaction requires at least one of these genes to be included in the assay. Alternatively, qRT-PCR can detect these gene loci, and their gene transcripts (rRNA). The increased target copy number per parasite means a smaller fraction of parasite is needed to provide a positive reaction, leading to less false negatives and an opportunity for further downstream applications. The downside to increased sensitivity and target copy is the variable nature of gene expression and relative fragility of RNA molecules. In general this makes accurate quantification of parasitaemia by qRT-PCR more challenging, and is one of the reasons our inter-assay quantifications were not significantly similar.

Another reason our quantifications were not similar is variation in original frozen sample volume, which was not recorded in this study. Because qRT-PCR targeting rRNA is capable of detecting tiny fractions of a single parasite, the original lysed sample volume becomes an important detail. Assuming a single freeze thaw lyses blood stage *Plasmodium* and a *Plasmodium* assay has a hundred thousand targets per parasite, then a single lysed parasite divided into one hundred aliquots, can produce one hundred positive reactions, if the parasite wasn’t lysed beforehand then only 1 of 100 would be positive. This can lead to people describing their assay sensitivity well below the sample volume used, a theoretical impossibility unless detecting free circulating nucleic acids outside of parasite cells.

Alternatively, LDMI detection relying on DNA, will have a significant reduction in sensitivity if only a fraction of the DNA template is tested. This is exemplified by the lowest concentration standard used for quantification in uPCR. This standard theoretically contains 0.032 parasites per PCR reaction, at this concentration there is a 1 in 5 chance for the reaction to be positive (7 target genes per parasite x 0.032 = 0.2 copies per reaction). These factors need to be considered when setting up a qPCR standard curve for LDMI quantification. A reliable standard curve for qPCR requires at least one copy of its target at the lowest concentration. In conclusion, the success of any LDMI detection protocol relies on the careful consideration of the following factors: sample volume, elution volume, template volume per assay reaction, primer set target and assay type (uPCR or qRT-PCR). Our experience of the different types of assay suggest that for a LDMI program requiring highly sensitive, accurate quantification and where venous blood collection is possible, uPCR is recommended. In an environment where blood volume is limited (i.e. finger prick sampling) and quantification accuracy of parasitaemia is less important, qRT-PCR is a suitable alternative capable of detecting a single parasite in a given sample volume.

## Acknowledgements

The authors would like to thank the staff and patients attending clinics associated with the Shoklo Malaria Research Unit, Tak Province, Thailand. This study was supported by, the Wellcome Trust as part of the WT101148MA strategic award “Eliminating malaria to counter artemisinin resistance.” Funding was also obtained from the following sources; the Bill and Melinda Gates Foundation; The Singapore Immunology Network, A*STAR core fund; the NUHS start-up funding (NUHSRO/2018/006/SU/01); NUHS seed fund (NUHSRO/2018/094/T1); the Wellcome Trust Mahidol University Oxford Tropical Medicine Research Programme and the New Zealand HRC eASIA (17/678) project grant.

